# Targeting study of HepG2 hepatoma cells in vitro by drug-loaded pectin-based nanoparticles

**DOI:** 10.1101/628818

**Authors:** Anil Shumroni, David Gupta

## Abstract

The biodegradable and biodegradable natural polysaccharide has always been used as a drug delivery system, and has the following advantages: It can prolong the biological half life of the drug and reduce the side effects of the drug. This experiment aimed to prepare a 5-fluorouracil (5-FU) nanoparticle (P-5-FU) drug-loading system based on pectin, and explored a large number of pectin-based nano drug-loading systems. The galactose residue is a natural target that targets human hepatoma cell HepG2. MTT assay was used to determine the proliferation inhibition effect of drug-loaded pectin-based nanoparticles on HepG2 and A549 cells. MTT assay showed that P-5-FU inhibited the proliferation of HepG2 cells in a dose-dependent manner, and the effect was stronger than 5-FU. P-5-FU also inhibited the proliferation of A549 cells in a dose-dependent manner, but there was no significant difference compared with 5-FU. High performance liquid chromatography (HPLC) on two kinds of cells loaded with drug-loaded nanoparticles the uptake and targeting were measured. The results of cell uptake showed that the uptake of P-5-FU by HepG2 cells was significantly higher than that of 5-FU, but there was no significant difference in the uptake of P-5-FU and 5-FU by A549 cells. There was no significant difference in the uptake of P-5-FU and 5-FU between the two cells after the galactose-saturated ASGPR binding site. The results indicate that pectin-based nano drug-loaded particles can specifically target highly expressed cells.

## Introduction

Cancer is the number one killer of humans. Hepatocellular carcinoma (HCC, hereinafter referred to as liver cancer) is the fifth most common tumor in the world [1]. According to statistics, more than 500,000 new cases of HCC are added each year, and more than 60,000 deaths are reported [1]. 5-Fluorouracil (5-FU) is one of the most widely used cytotoxic and anti-tumor drugs, and shows good therapeutic effects on cancers such as colon cancer, breast cancer, gastrointestinal cancer, and liver cancer. It is also accompanied by strong side effects such as myelosuppression, gastrointestinal reactions, leukopenia, thrombocytopenia, etc. This may be due to higher blood levels and non-specific systemic distribution of anticancer drugs [2] This severely limits its clinical application. In addition, the absorption of the drug in the gastrointestinal tract after oral administration is non-uniform, which is caused by the metabolism of dihydropyrimidine dehydrogenase or uracil reductase [3]. Therefore, in order to avoid these shortcomings, drug delivery systems such as: nanoparticles, liposomes, microemulsions, etc. came into being [4–7].

Targeted drug delivery (TDD) refers to the way in which drugs are concentrated in target tissues, target organs, and target cells through local or systemic blood circulation, and can be divided into active targeting and passive targeting. [8]. Nanoparticles, liposomes and micelles can be passively targeted to tumor tissue by the EPR (Enhanced Permeability and Retention) effect, and some enzymes and receptors in tumor cells can also be utilized by grafting ligands on nanocarriers. The high level of expression achieves active targeting [8–10]^1-21^.

The asialoglycoprotein receptor (ASGPR) is a transmembrane protein that is mainly present on the surface of mammalian liver parenchyma cells and specifically recognizes and binds to glycoproteins with galactose residues at the ends of the molecules. Lysosomes in hepatocytes are metabolized [11]. Each liver cell contains approximately 2 million ASGPR binding sites [12]. Therefore, researchers began to study drug delivery systems targeting asialoglycoprotein receptors, hoping that drug delivery systems with galactose residues can deliver antitumor drugs to tumor tissue through blood circulation, thereby increasing anticancer activity. Local concentration of the drug [13–14]. In recent years, researchers have focused on the study of natural polymer nanoparticles due to their good biocompatibility, non-toxicity, biodegradability and adjustable controlled release properties [15–16]. Pectin is one of the most widely used natural polysaccharides with a large amount of galactose residues, and galactose can bind to ASGPR, thus achieving active targeting of hepatocytes. In the early stage of our research group, we have combined the results of the drug-loaded nanoparticles, and studied the particle size, morphology, drug loading and in vitro release characteristics [17].

The purpose of this study was to evaluate the cytotoxicity and cellular uptake of drug-loaded nanoparticles in vitro. Studies have found that some tumor cells such as Caco-2, HT-29 [18] colon cancer cells and HepG2 70 Vol. 27, No. 1 Wang Yanmei, etc.: Targeted research on HepG2 hepatoma cells loaded with pectin-based nanoparticles in vitro [19] The surface also has the expression of asialoglycoprotein receptor, and A549 lung cancer cells are typical cells without ASGPR expression. Therefore, A549 lung cancer cells were used as controls for HepG2 liver cancer cells for in vitro cytotoxicity. Evaluation of cellular uptake and targeting.

## Materials and Method

HP-1100 High Performance Liquid Chromatograph (Agilent, USA) 5-fluorouracil (5-FU), pectin (Sigma, P9135, Mw = 1. 18 × 105 g / mol, multi-angle laser light scattering measurement), carbonic acid Sodium hydrogen (Shanghai Sinopharm), calcium hydroxide (Tianjin Komiou Co., Ltd.). MTT (Amresco, USA), acetonitrile (Tianjin Komio Co., Ltd.), DMEM medium (Hyclone, USA), fetal bovine serum (Gibco, USA), HepG2 human liver cancer cell line (Shanghai Institute of Life Sciences), A549 Lung cancer cell line (courtesy of Chen Linxi, College of Pharmacy and Life Sciences, University of South China), the rest of the reagents were of analytical grade.

Preparation, characterization, drug loading, encapsulation efficiency and in vitro release of 5-FU pectin-based nanoparticles were performed using our previous method [17].

Take HepG2 liver cancer cells and A549 lung cancer cells in logarithmic growth phase, aspirate the culture solution, and use 0. 25% trypsinization, stop digestion with serum-containing medium, prepare cell suspension, adjust cell density to 5 × 104 /mL, gently blow and inoculate in 96-well plate, 200 μL per well, set at 5 Incubate culture at % CO2, 37 °C, and aspirate the culture solution after 24 h, and add the culture medium containing 5-FU and P-5-FU, respectively, and set 5, 1. 7, 0. 56, 0. 19, 0. 0063, 0. 7 concentrations such as 0021, 0. 0007 mmol / L and the control group, 3 duplicate wells in each group. Continue to incubate in 5% CO2, 37 °C. After 48 h, add 5 μL of 5 mg /mL MTT per well. Incubate in the incubator for 4 h, then discard the supernatant and add 150 μL of DMSO to each well. After shaking for 10 min, the crystals were fully dissolved and mixed. The absorbance of each well at a wavelength of 570 nm was measured by a microplate reader. The inhibition rate of each group was calculated according to the formula: Inhibition rate % = (OD value of the control group - administration) Group OD value) / control group OD value × 100%, and use SPSS 18. 0 Calculate IC50 values of 5-FU and P-5-FU on HepG2 hepatoma cells and A549 lung cancer cells.

Take HepG2 liver cancer cells and A549 lung cancer cells in logarithmic growth phase, aspirate the culture solution, and use 0. 25% trypsinization, stop the digestion with serum-containing medium, make a cell suspension and adjust the cell density, gently blow it, and inoculate it in a 6-well plate at a density of 1 × 106 /well, each containing culture medium 2 mL, placed in an incubator culture at 5% CO2, 37 °C, and changed every 2 days until the cells covered the bottom of the well. Replace the fresh medium 24 hours before the start of the experiment. 1 h before administration, the culture solution was aspirated, gently washed twice with normal temperature PBS, and then 1 mL of HBSS was added (additional group added 800 mmol / L galactose to observe galactose on HepG2he A549 cells to P -5-FU uptake effect), incubate at 37 °C for 120 min. Aspirate the supernatant and add 0. 5 mM/L of 5-FU and P-5-FU of HBSS, at 120 min, respectively (with a blank control group), aspirate the drug-containing supernatant, wash 3 times with pre-cooled HBSS, then add 200 μL deionized water, placed in an −80 degree refrigerator, set with 3 duplicate holes.

## Results and Discussion

Preparation methods, characterization parameters, drug loading, encapsulation efficiency and in vitro release of drug-loaded nanoparticles refer to the previous method of the research group [17]. The encapsulation rate has reached 37. 7%, drug loading 28. 14% and at flat pH = 7. 4. The neutral solution of 4 showed obvious controlled release characteristics. Figure 1 is a transmission electron micrograph of pectin-based nanoparticles. The results of TEM electron microscopy show that the average particle size of pectin-based nanoparticles is between 40 and 80 nm.

**Figure 1.**
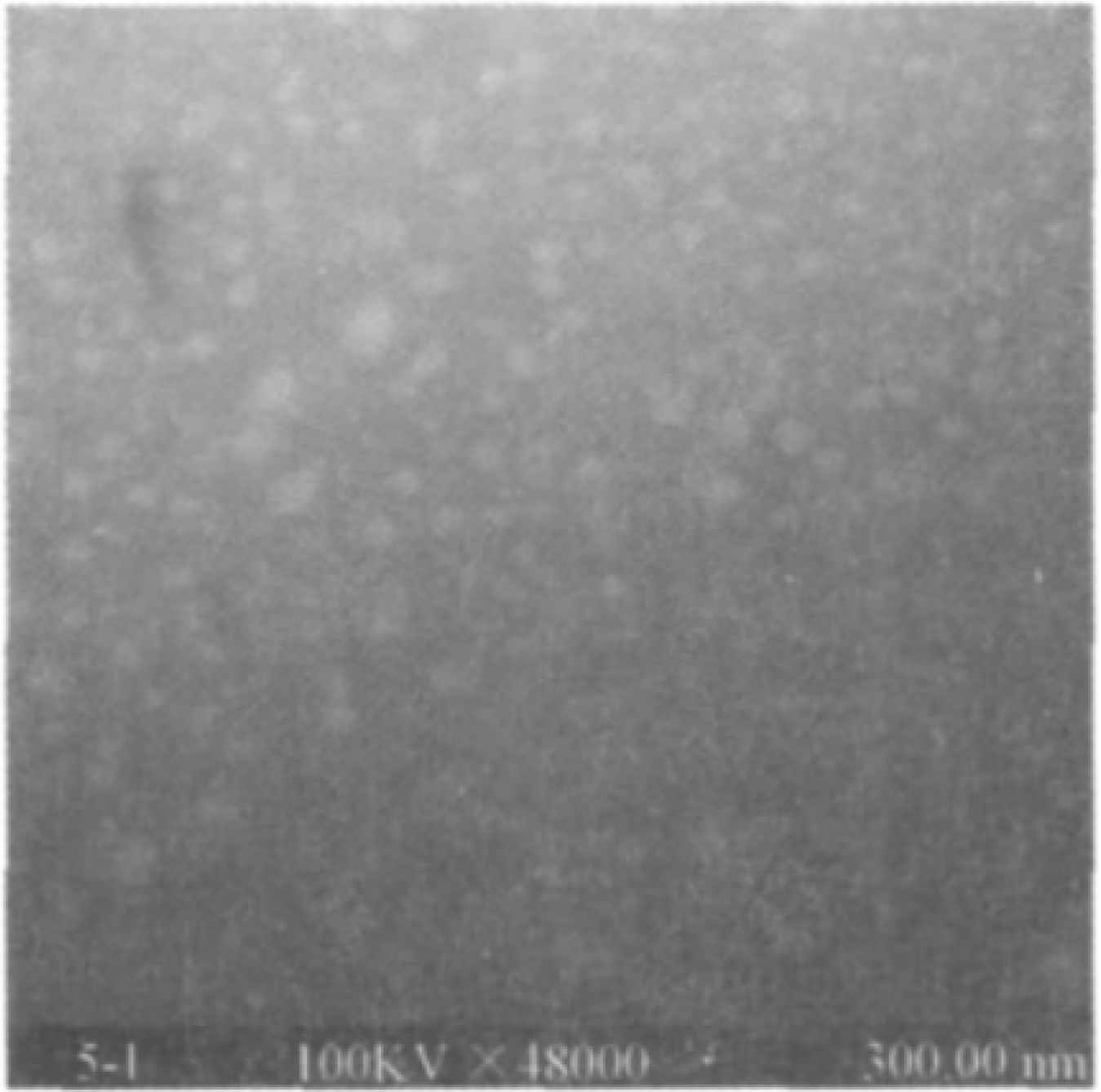
TEM analysis

In this study, MTT assay was used to investigate the inhibitory effects of 5-FU, P-5-FU and blank nanoparticles on the proliferation of HepG2 hepatoma cells and A549 lung cancer cells. The concentration of blank nanoparticles was the same as that of drug-loaded nanoparticles. Different concentrations of blank nanoparticles did not inhibit the two cancer cells. The highest inhibitory rate of the highest concentration of blank nanoparticles on HepG2 and A549 cells was 4. 07% and 10. 07%, Figure 2 shows the survival rate of the two cells treated with high concentration of blank nanoparticles for 48 h, indicating that the prepared pectin-based nanoparticles have good biocompatibility.

**Figure 2.**
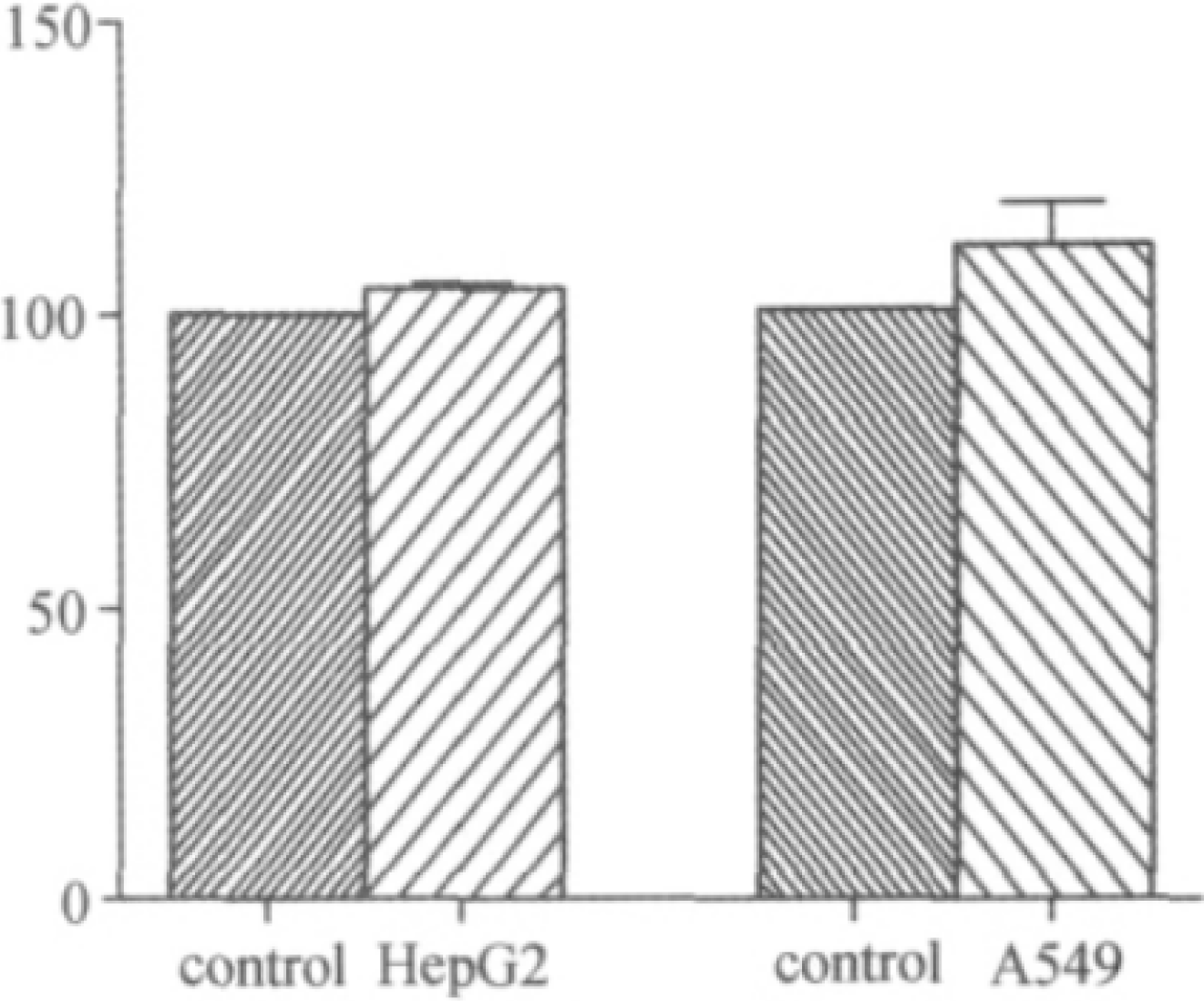
Cytotoxicity analysis

3 indicates the inhibitory effect of 5-FU and P-5-FU on the proliferation of HepG2 and A549 tumor cell lines. Figure 3A shows that P-5-FU has a significant inhibitory effect on the proliferation of HepG2 cells (p < 0.05) in a dose-dependent manner, which is significantly enhanced compared with 5-FU; whereas Figure 3B shows that P-5-FU and 5 There was no significant difference in the inhibitory effect of -FU on proliferation of A549 cells (p > 0.05). As can be seen from Table 1 and Figure 3, the IC50 value of P-5-FU was significantly lower than that of free 5-FU for HepG2 cells, and the IC50 value of P-5-FU for A549 cells. There was no significant difference between the IC50 values of free 5-FU.

**Figure 3.**
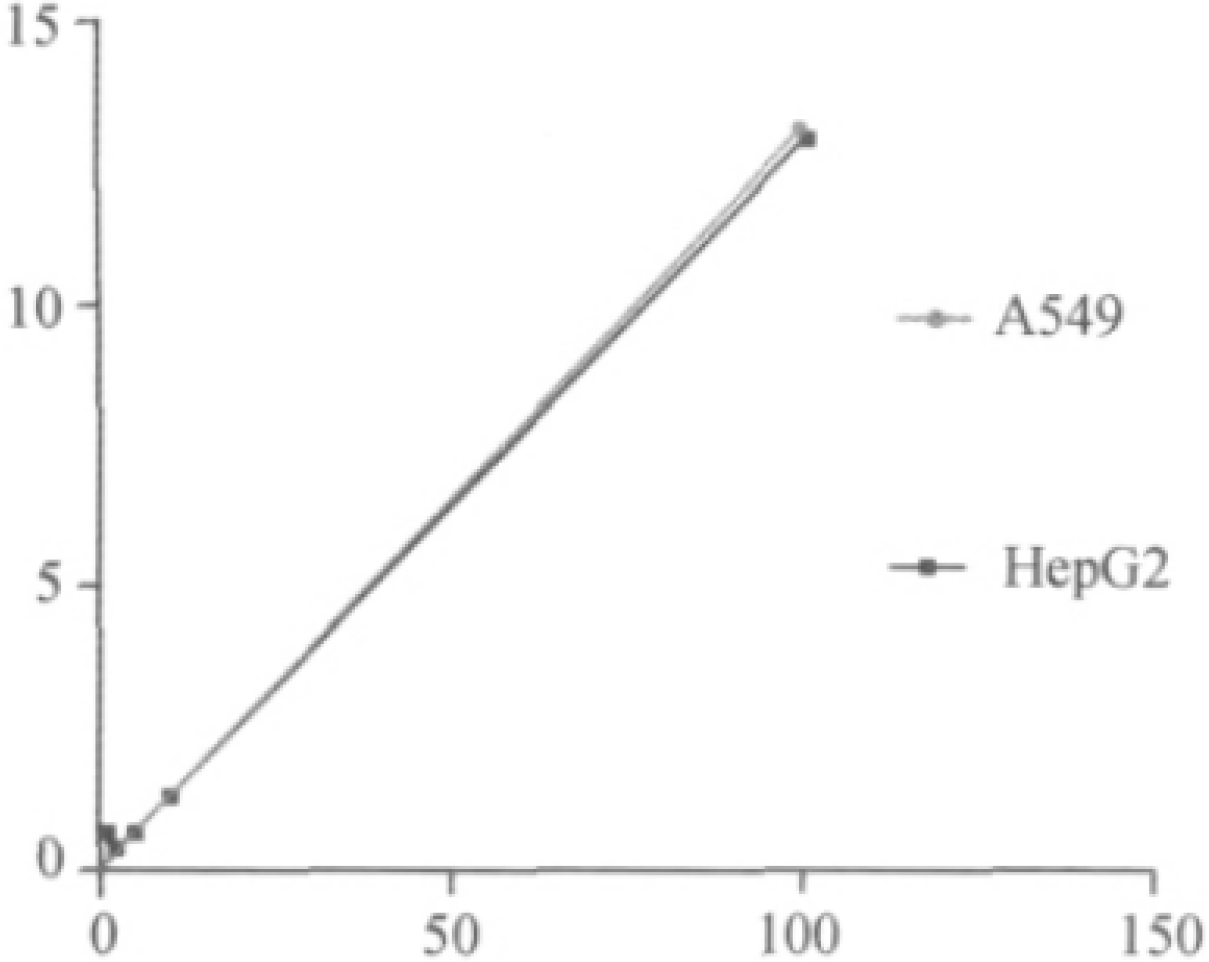
Releasing kinetics

**Figure 4.**
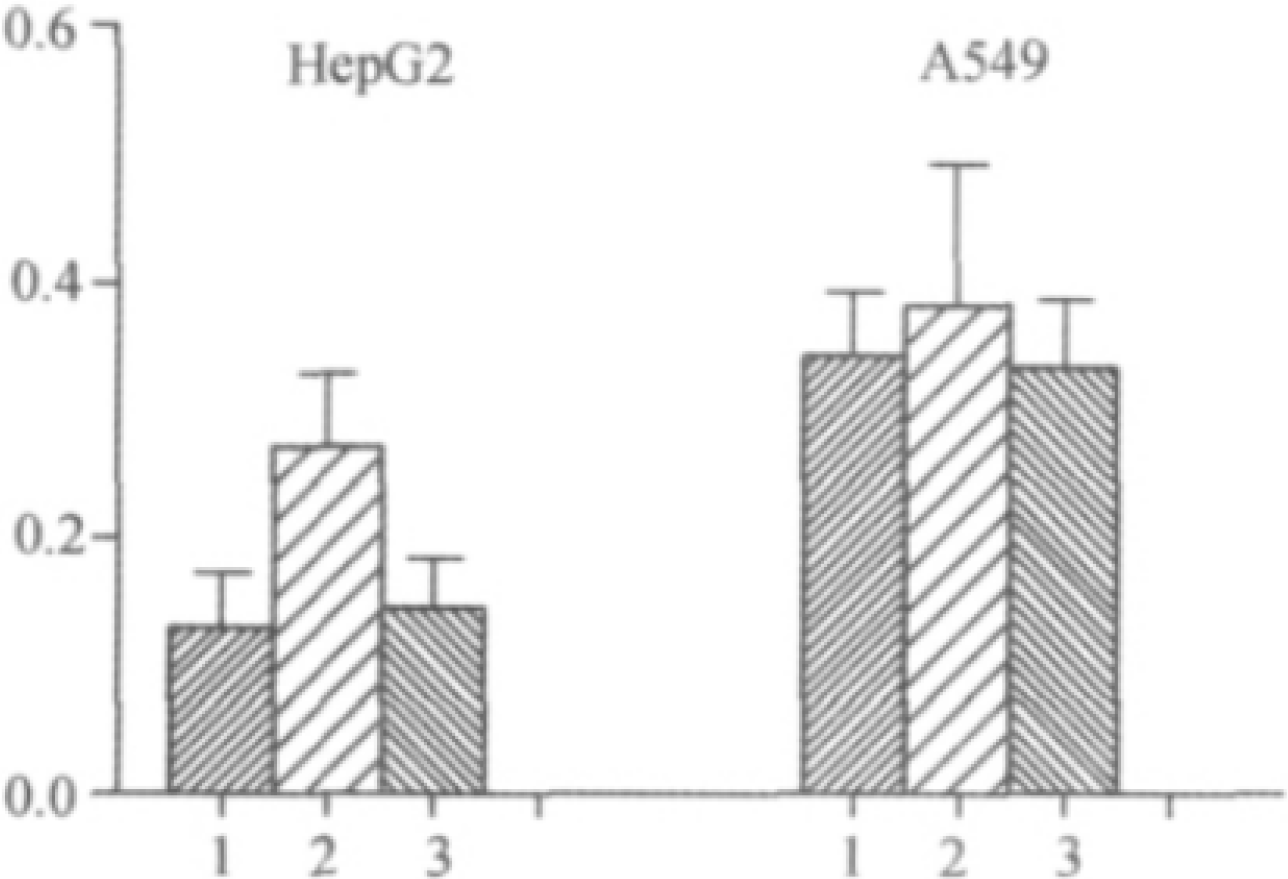
Cytotoxicity comparison between different cell types

The 5-Fu samples in the low, medium and high concentrations of 5-Fu in HepG2 and A549 blank cell fluids were divided by the chromatographic peak area obtained after extraction and the chromatographic peak area obtained by direct extraction without extraction. 85%, see Table 2.

The HPLC method was used to determine the uptake of 5-FU and P-5-FU by two cells. Galactose can bind to ASGPR and competitively inhibit the binding of ASGPR to galactose-containing particles [20]. In order to verify that the difference in binding between P-5-FU and 5-FU and HepG2 cells is related to ASGPR-mediated, galactose is pre-added to HepG2 cells or A549 cells in a 6-well plate, and the binding site of ASGPR on the surface of pre-saturated cells is pre-saturated. Point, then add P-5-FU (GP-5-FU) and 5-FU. As shown in Figure 5. The uptake of P5-FU by HepG2 is 5-FU. At 1 time, the binding of P-5-FU to HepG2 cells was significantly decreased after adding galactose, and there was no difference in the uptake of 5-FU. There was no significant difference in the uptake of P-5-FU and 5-FU between A549 cells and the addition of galactose had no effect on the binding of P-5-FU and 5-FU to A549 cells. These results indicate that the increase in binding of P-5-FU to HepG2 cells is achieved by ASGPR-mediated pathway.

It is suggested that P-5-FU may have a targeting effect on tumor cells with high expression of ASGPR. The possible mechanism is that the galactosyl group of 5-FU galactosidase can specifically recognize the ASGPR receptor on the surface of tumor cells and mediate endocytosis. And accelerate the uptake of tumor cells.

## Conclusion

In recent years, ASGPR-mediated targeting is a hotspot in the research of liver-targeted drug delivery systems. The use of natural pectin-based 5-FU nanoparticles [17], using the pectin-based nano drug-loading system itself with a large number of galactose residues as a natural target to achieve active targeting, verified its HepG2 cells Proliferation inhibition rate of A549 cells (including ASGPR receptors) and A549 cells (without ASGPR receptors) and uptake of pecto-based nano drug-loading systems by two cell lines. Experiments showed that the binding amount of P-5-FU to HepG2 cells was significantly higher than that of 5-FU; however, the binding amount of P-5-FU was not different from that of A549 cells; while ASGPR on the surface of HepG2 cells was presaturated by galactose, P-5-FU The binding amount to HepG2 cells was significantly reduced, and the binding amount to 5-FU was basically the same. It was confirmed that the galactose residue exposed on the surface of the nanoparticles in P-5-FU can be specifically recognized by ASGPR on the surface of HepG2 cells, and the pectin-loaded nanoparticles have the characteristics of actively targeting hepatoma cells. It indicates that pectin-based nanoparticles have a good application prospect as a drug delivery system targeting liver diseases.

